# Hippocampal-Prefrontal Interactions during Spatial Decision-Making

**DOI:** 10.1101/2020.06.24.168732

**Authors:** Lucas CS Tavares, Adriano BL Tort

## Abstract

The hippocampus has been linked to memory encoding and spatial navigation, while the prefrontal cortex is associated with cognitive functions such as decision-making. These regions are hypothesized to communicate in tasks that demand both spatial navigation and decision-making processes. However, the electrophysiological signatures underlying this communication remain to be better elucidated. To investigate the dynamics of the hippocampal-prefrontal interactions, we have analyzed their local field potentials and spiking activity recorded from rats performing an odor-cued spatial alternation task in an 8-shaped maze. We found that the phase coherence of theta peaked around the choice point area of the maze. Moreover, Granger causality revealed a hippocampus->prefrontal cortex directionality of information flow at theta frequency, peaking at starting areas of the maze, and on the reverse direction at delta frequency, peaking near the turn onset. Additionally, the patterns of phase-amplitude cross-frequency coupling within and between the regions also showed spatial selectivity, and a new method revealed that hippocampal theta and prefrontal delta modulated not only gamma amplitude but also inter-regional gamma synchrony. Lastly, we found that the theta rhythm dynamically modulated neurons in both regions, with the highest modulation at the choice area; interestingly, prefrontal cortex neurons were more strongly modulated by the hippocampal theta rhythm than by their local field rhythm. In all, our results reveal maximum electrophysiological interactions between the hippocampus and the prefrontal cortex near the decision-making period of the spatial alternation task. These results corroborate the hypothesis that a dynamic interplay between these regions takes place during spatial decisions.

## Introduction

The hippocampus has been extensively linked with memory formation (Squire et al., 1992) and spatial navigation (Moser et al., 2008; O’Keefe, 1991). Regarding the latter, the hippocampus is believed to be part of a system for assessing positional information and the surrounding context, which allows subjects to navigate across space. On the other hand, the prefrontal cortex plays a role in selecting context-specific memories among competing, non-appropriate ones, acting as a control mechanism for the functional discrimination of information (Dobbins et al., 2002; Moscovitch, 1992; Preston and Eichenbaum, 2013; Szczepanski and Knight, 2014). The interplay between these regions is thus expected to take place in tasks or behaviors that demand both navigational data and choice-related memory discrimination.

Several studies have demonstrated a link between animal behavior and connectivity measures (Fell et al., 2001; Varela et al., 2001; Womelsdorf et al., 2007). Electrophysiological interactions between distant brain regions are usually mediated by low-frequency oscillations (Daume et al., 2017; A. von Stein and Sarnthein, 2000). Among them, theta oscillations (6-10 Hz) can be detected at the single-cell level (Klausberger et al., 2003) as well as at the population level in a variety of brain regions, such as the hippocampus (Buzsáki, 2002), the neocortex (Sirota et al., 2008), thalamic and subthalamic nuclei (Kirk et al., 1996; McNaughton et al., 1995). The hippocampal theta cycle modulates spiking activity from distal neuronal populations (Buzsáki, 2006; Wang, 2010), while theta phase coherence has been suggested as a potential mechanism for the synchronization of the prefrontal-hippocampal network during cognitive tasks (Benchenane et al., 2010; Jones and Wilson, 2005a).

Recent reviews highlight the role of hippocampal-prefrontal interactions across different cognitive domains, such as goal-directed behavior (Womelsdorf et al., 2010), emotion (Jin and Maren, 2015), context-guided memory (Place et al., 2016), episodic memory (Eichenbaum, 2017), decision-making (Tamura et al., 2017) and spatial learning (Maharjan et al., 2018). Here, we sought to revisit and further characterize the electrophysiological signatures of hippocampal-prefrontal interactions during spatial decision. To that end, we have analyzed local field potentials (LFPs) and spiking activity previously recorded from the hippocampus and prefrontal cortex of rats performing an odor-cued alternation task in an 8-shaped maze (Fujisawa et al., 2008).

## Results

We analyzed 13 sessions from 3 Long Evans rats performing a spatial task which required them to leave the start area, cross the center arm of the maze, and then choose between the left or right goal arm for reward (Figure 1A). The correct arm and particular reward (300 mg of cheese or chocolate) were instructed by a matched odor sample (i.e., cheese or chocolate scent) delivered at the start area upon a nose poke. Rats were proficient at this task (>85% correct choices) before surgery for implanting multi-shank multi-contact silicon probes in the CA1 region of the hippocampus and medial prefrontal cortex (mPFC). For the LFP analyses that follow, we have focused on one channel for each region selected based on the highest theta/delta ratio, but the results hold with other channel choices given the high redundancy among signals from the same region.

**Figure 1.**
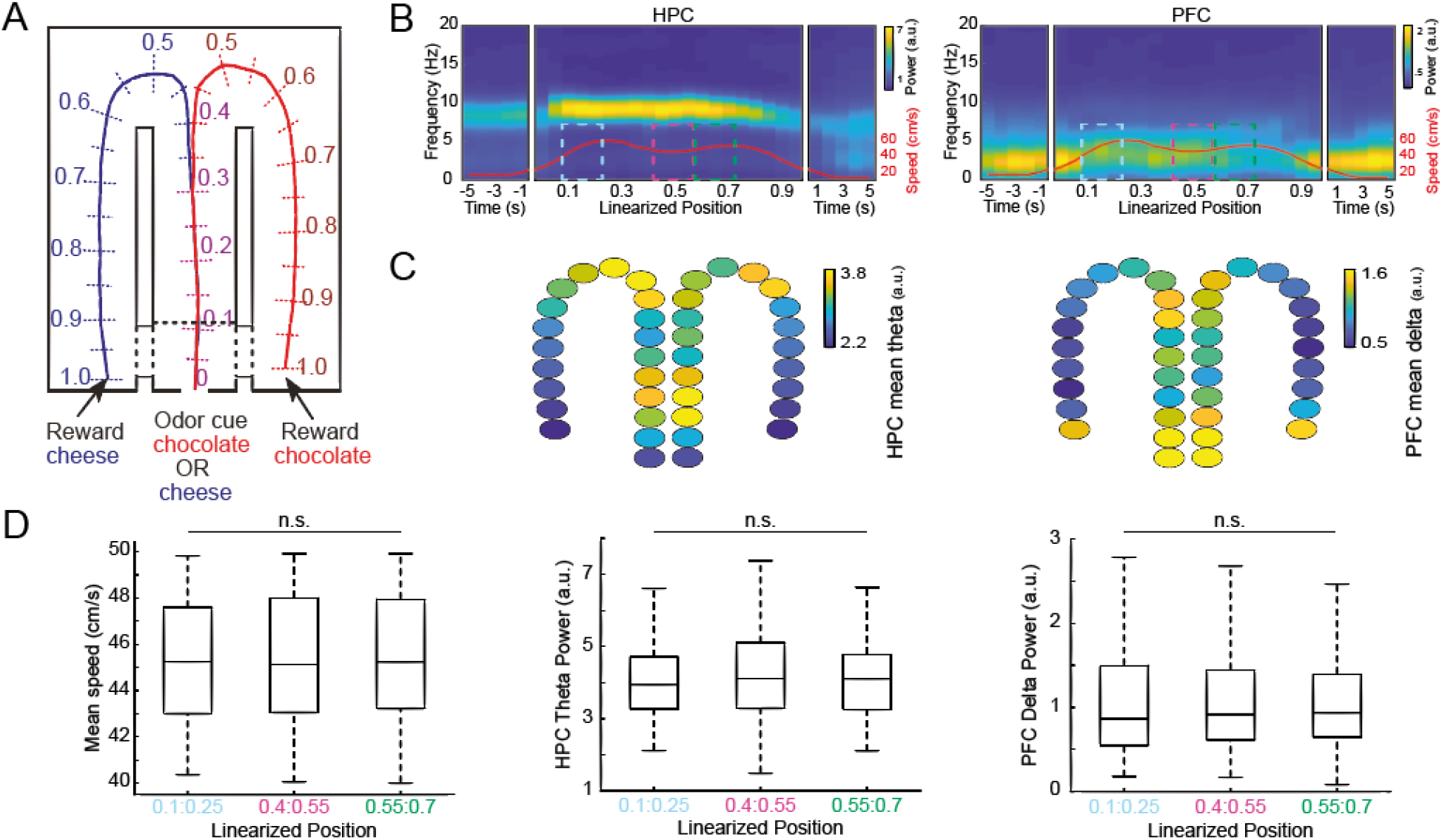
HPC and PFC oscillatory power content varies upon maze runs but does not differentiate between pre- and post-decision periods when controlling for speed. (A) Schematic representation of the 8-shaped maze where animals performed an odor-cued spatial decision task (adapted from Fujisawa et al., 2009; see Methods). The lines denote the mean trajectory during left (blue) and right (red) runs, which were converted into a linearized position ranging from 0 to 1. (B) Average spatial spectrograms for LFPs from the hippocampus (HPC) and prefrontal cortex (PFC) (20 spatial bins; n = 13 sessions). The red line denotes the mean locomotion speed. The dashed rectangles demark spatial bins analyzed in D. HPC exhibits prominent theta activity through most of the trajectory, while PFC power peaks at delta frequency and is highest during low speeds. (C) Average HPC theta power and PFC delta power across spatial bins. Notice a power maximum in HPC theta before the maze bifurcation but also at the turn. Higher values of PFC delta power are present in regions of lower speed. (D) Left: Boxplot distributions of the mean locomotion speed for the subsets of trials analyzed in the middle and right panels (n=146 trials for each maze subregion). Middle and right: CA1 theta power and PFC delta power in spatial locations before and after the maze bifurcation for the speed-controlled trials. n.s.: not significant (one-way ANOVA).

We started by investigating the LFP power content while animals executed the task. To better visualize the relationship (or lack thereof) of oscillatory activity with the trial space, we divided the maze into 20 equally-spaced bins, referred to as linearized positions from 0 to 1, and computed the average spectrogram for each spatial bin. In Figure 1B, we plot the average spectral content for each region as a function of linearized position, along with five 1-second time bins before and after the trial executions (n = 13 sessions).

The results show a prominent theta-band oscillation centered around 9 Hz in the CA1 region during maze runs, which had otherwise low amplitude at moments before and after the trial (Figure 1B left). Since animals have long periods of immobility during inter-trial intervals, this power dynamics is in agreement with the vast literature correlating hippocampal theta oscillations with locomotion (Whishaw and Vanderwolf, 1973; McFarland et al., 1975; Sławińska and Kasicki, 1998; Buzsáki, 2002).

For the mPFC, the most prominent oscillation occurred in the delta band and was highest during periods of low locomotion before and after the trial, as well as in regions near the curve, where animals slowed down locomotion velocity (Figure 1B right). Of note, although various behaviors have been associated with delta oscillations in the mPFC (Karalis et al., 2016), we highly suspect that this rhythm corresponds to respiration-entrained LFP oscillations, which have recently been shown to be prominent in prefrontal regions (Tort et al., 2018a; Lockmann and Tort, 2018) especially during immobility (Biskamp et al., 2017; Zhong et al., 2017). At any event, since respiration was not concomitantly recorded, we cannot draw conclusive inference in this regard (but see Discussion).

In Figure 1C, we show schematic representations of the mean CA1 theta power (left) and mean mPFC delta power (right) across 20 space bins, separately for left and right runs. These plots reveal similar spatial power distributions in the center arm irrespective of subsequent arm choice, and virtually symmetric power values between the left and right arms. Moreover, they also show the already mentioned maximum mPFC delta power at maze start and end, as well as the high CA1 theta power at the center arm. Since the analyzed bins encompass only the trial execution space, locomotion is present in all of them and cannot be accounting for the power changes in a binary way. Nevertheless, the changes in oscillatory content matched well the changes in locomotion speed (red curve in Figure 1B panels).

In addition to the clear influence of speed, we next sought to determine if any power variation could be related to the (presumed) differences in cognitive demands during trial execution. To that end, we analyzed LFPs associated with three maze subregions while controlling for locomotion speed. That is, for each maze subregion, we selected a subset of trials such as to obtained similar distributions of speed values among the three subregions (one-way ANOVA, F(2,435)=0.16, p=0.85; Figure 1D left). The selected spatial subregions included either data before (“choice point”), during (“turn start”), or after (“turn end”) the bifurcation of the mean locomotion trajectory, that is, before and after the decision point. Under this scenario, we found no significant differences among the spatial subregions in CA1 theta power (F(2,435)=0.51, p=0.60; Figure 1D middle) nor in mPFC delta power (F(2,435)=0.05, p=0.95; Figure 1D right). Therefore, the variations in oscillatory power in this spatial-alternation task could not be explained by differences in cognitive load across task execution, and are likely to be entirely accounted for by changes in locomotion speed.

We next moved on to analyze inter-regional LFP interactions. We started by computing phase coherence across the frequency spectrum, which measures the stability of phase differences between two signals at the same frequency. We estimated phase coherence throughout trial execution using the same time and spatial bins as described above for the analysis of oscillatory power (see Methods). Phase coherence between the two regions was most prominent at the theta frequency band (Figure 2A). Interestingly, CA1-mPFC theta coherence peaked at the middle of the center arm, thus before maze bifurcation (Figure 2B), in an area thought to be associated with choice due to its proximity to the spatial decision commitment, as inferred by the separation between the mean locomotion trajectories for left and right choices (see Figure 1A). The spatial distribution of inter-regional theta coherence was similar for left and right runs (Figure 2B). Interestingly, we found that the spatial profile of theta coherence differed from the spatial profile of theta power. For instance, Figure 2C shows theta power and coherence for spatial locations before and after maze bifurcation. Notice that while theta power was not different in the two locations (mean band power: t(12)=1.18, p = 0.26; peak power: t(12)=-0.032, p = 0.97; paired t-tests), theta coherence was statistically significantly higher before the curve (mean band coherence: t(12)=5.46, p<0.001; peak coherence: t(12)=6.23, p<0.0001; paired t-tests; Figure 2C).

**Figure 2.**
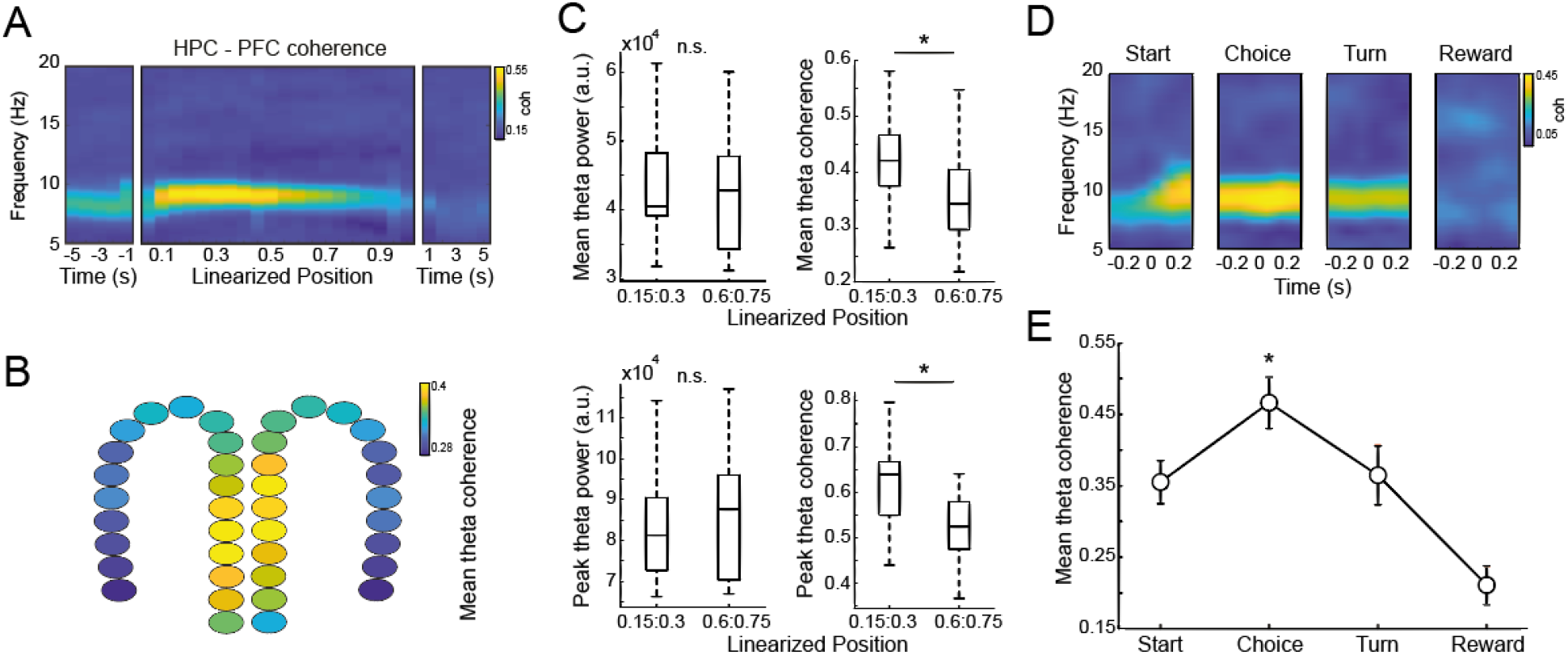
HPC-PFC theta coherence peaks during spatial decision. (A) Average HPC-PFC coherogram (n = 13 sessions). (B) Schematic representation of HPC-PFC theta coherence across spatial bins. Theta coherence peaks before the maze bifurcation. (C) Boxplots of theta power and theta coherence for spatial locations before and after the turn. The top and bottom panels show band average and peak values, respectively. Notice statistically higher theta coherence before maze bifurcation, along with no difference in theta power. *p<0.001, paired t-test. (D) Time-resolved HPC-PFC coherogram centered at task events (see Methods). Notice the highest theta coherence at the choice period. (E) HPC-PFC theta coherence per task event (*p<0.0001, repeated measures ANOVA).

Since the accuracy of coherence estimates depends on the length of the analyzed time (i.e., the number of analyzed cycles), it has an intrinsic tradeoff with spatial resolution (i.e., the higher the spatial resolution, the less time the animal spends in a given spatial bin). We next employed a second approach for estimating phase coherence which loses spatial precision in favor of longer and consistently-sized time windows. In specific, in this so-called “task-event approach”, we computed coherence estimates triggered by time-stamps around events of interest during trial execution, such as when the animals reached a particular position of the maze (see Methods). Using this approach, we once again observed a peak in theta phase coherence around the choice point event (Figure 2D), which was statistically significantly higher than theta coherence in other task events (F(3,36)=24.54, p<0.0001, repeated measures ANOVA; Figure 2E). Therefore, both the spatial- and time-triggered estimates support the conclusion of high mPFC-CA1 theta coherence during spatial decision-making.

The phase coherence analysis performed above does not quantify directional relations between the regions. To address the directionality of information flow, we next used Granger causality (GC), a metric that calculates the level of prediction a time series has over another. Given that we were interested in oscillatory interactions, here we computed Granger causality in the spectral domain (Geweke, 1982, 1984). This metric has been previously applied in several fields of neuroscience for assessing directed connectivity at specific frequencies (Ferri et al., 2007; Liao et al., 2010; Bressler and Seth, 2011; Seth et al., 2015). Interestingly, the results revealed a prominent theta peak in the GC spectrum in the hippocampus to prefrontal direction, and a causality peak at delta frequency in the reverse direction (Figure 3A top). Perhaps most strikingly, we found that the magnitude of this functional directed connectivity significantly varied across trial execution. Namely, when employing the task-event approach (Figure 3A bottom), the highest CA1->mPFC theta causality occurred in the choice point task event (F(3,36)=66.71, p<0.00001; repeated measures ANOVA), while mPFC->CA1 delta causality peaked at the turn event (F(3,36)=4.52, p<0.01; repeated measures ANOVA). Interestingly, two-way ANOVA revealed not only higher Granger causality for the CA1->mPFC direction (F(1,96)=108.73, p<10^-16^), but also an interaction effect (F(3,96)=16.23, p<10^-7^), showing that the changes in causality across task execution depend on flow direction. Consistent results were obtained when characterizing GC throughout space (using ten equally spaced position bins; see Methods): the highest CA1->mPFC GC occurred in initial areas of the center arm, except for the initial box, and highest delta mPFC->CA1 GC occurred at maze bifurcation (Figure 3B,C). Thus, the LFP oscillatory causality between the regions occurs at direction-specific frequencies and changes dynamically while animals execute the task.

**Figure 3.**
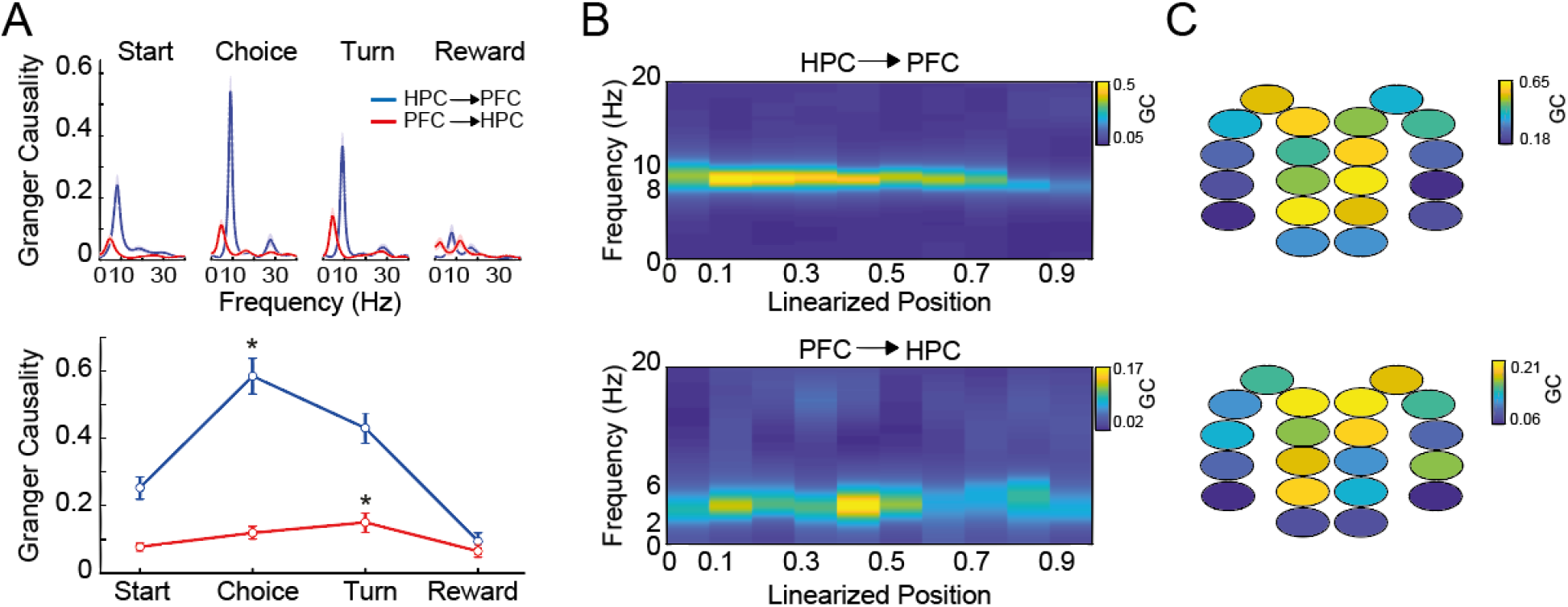
HPC->PFC and PFC->HPC causality peak at different frequencies and maze regions. (A) Top: spectral Granger causality during task events (average over 13 sessions). In the hippocampus-prefrontal direction (blue), causality peaks at theta and is highest at the choice point, while the prefrontal-hippocampus direction (red) peaks at delta and is highest at the turn. Bottom: HPC->PFC theta (blue) and PFC->HPC delta (red) Granger causality for each task event (*p<0.0001, repeated measures ANOVA). (B) Average spatial Grangerograms (see Methods). (C) Schematic representations of mean HPC=>PFC theta and PFC->HPC delta (bottom) Granger causality through 10 spatial bins. Note the peak theta causality in early areas of the maze and peak delta causality at turn onset.

In addition to characterizing power content and functional connectivity at the same frequencies, we also investigated the LFP interactions among oscillations of different frequencies, within and between the regions. These are commonly called cross-frequency coupling (CFC) or comodulation (Jensen and Colgin, 2007; Scheffer-Teixeira and Tort, 2018). Comodulation can happen between distinct properties of the oscillations, such as phase, amplitude, and frequency (Jensen and Colgin, 2007; Scheffer-Teixeira and Tort, 2018). Here we focus on phase-amplitude coupling (PAC), a relationship that has been hypothesized as a mechanism for functional communication between local and global circuits (Canolty and Knight, 2010). Within the CA1 region, we observed the presence of theta-gamma PAC, with amplitude frequencies varying from 40 to 80 Hz (Figure 4A), as largely reported (Colgin, 2015; Scheffer-Teixeira et al., 2012; Scheffer-Teixeira and Tort, 2017). On the other hand, and consistent with recent observations (Zhong et al., 2017; Andino-Pavlovsky et al., 2017), the mPFC exhibited local PAC between LFP phases near delta and faster gamma oscillations from 80 to 100 Hz (Figure 4A). Inter-regionally, we found no evidence of PAC between the amplitude of PFC frequencies and the phase of CA1 frequencies, thus similar to recent reports (Zhang et al., 2016; but see Sirota et al., 2008). Instead, comodulation was apparent in the opposite direction: the phase of PFC theta modulated the amplitude of CA1 gamma (Figure 4A; see also Zhang et al., 2016). It should be noted, however, that such cross-regional coupling is mathematically expected given the high phase coherence at theta between CA1 and mPFC, along with the high theta-gamma coupling in CA1.

**Figure 4.**
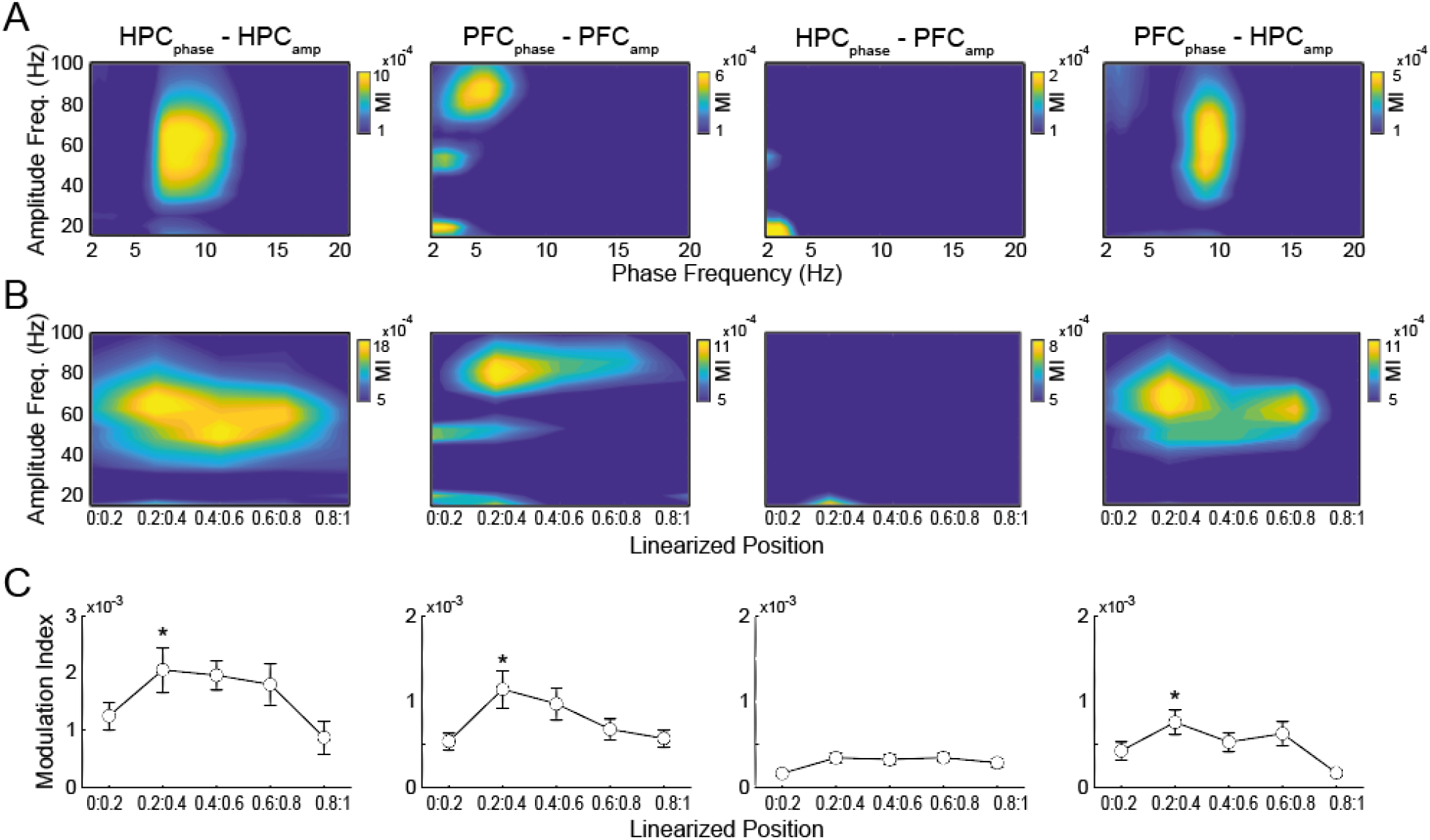
Spatial dynamics of hippocampal-prefrontal cross-frequency coupling. (A) Average whole-trial comodulograms (n = 13 sessions). The title states the region from where the phase (first) and amplitude (second) time series were obtained. In the intra-regional cases, CA1 theta phase modulates 40 to 80 Hz gamma, while the mPFC delta phase modulates faster gamma oscillations at 80 to 100 Hz. In the inter-regional cases, mPFC amplitude has no meaningful coupling to CA1 phase, while CA1 gamma couples to mPFC theta phase. (B) Average spatial comodulograms for amplitude frequencies (see Methods). CA1 and mPFC amplitude-based comodulograms were phase averaged in relation to theta and delta, respectively. (C) Modulation index across space for the relevant frequency pairs in A and B (mean over 13 sessions, error bars denote SEM; *p<0.01, repeated measures ANOVA).

We next computed the spatial dynamics of PAC for the relevant frequency ranges mentioned above. Theta-gamma PAC within CA1 was lower for maze areas of low locomotion, and otherwise high when animals crossed the maze (Figure 4B,C; F(4, 48)=8.11, p<0.001, repeated measures ANOVA). mPFC delta-gamma coupling exhibited a peak near the maze bifurcation (Figure 4B,C; F(4, 48)=5.42, p=0.001, repeated measures ANOVA). Two-way ANOVA confirmed higher levels for CA1 theta-gamma than PFC delta-gamma coupling (F(1, 120)=26.76, p<10^-15^) and the effect of space (F(4, 120)=4.56, p=0.0018), but showed no interaction effect (F(4, 120)=0.84, p=0.505). Once again, CA1 phase-mPFC amplitude had low comodulation levels, while the spatial dynamics of theta-gamma coupling between the mPFC and CA1 was similar to the dynamics for coupling within CA1, though at lower magnitude (Figure 4B,C; F(1,120)=52.96, p<10^-20^ for the phase region factor, two-way ANOVA).

We then moved on to investigate for possible relationships between CFC and communication through coherence (CTC). Here CTC was assessed as the long-distance (i.e., CA1-mPFC) synchrony at gamma frequencies, measured by the phase-locking value (PLV; Lachaux et al., 1999; Varela et al., 2001). To study the influence of CFC over CTC, we used a new screening method (González et al., 2020) that measures if the CA1-mPFC PLV for different gamma sub-bands (20 Hz bandwidths, see Methods) depends on the phase of slower oscillations recorded in one of the regions. This analysis revealed that the predominant slow oscillation in each region modulated long-range synchrony of different gamma sub-bands (Figure 5). Namely, the phase of hippocampal theta waves modulated inter-regional synchrony at 30-40 Hz, while the phase of prefrontal delta waves modulated synchrony at the 50-60 Hz gamma sub-band (p<0.05, surrogate analysis; Figure 5). These results thus suggest that different frequency channels may be used in the interaction between CA1 and mPFC depending on flow direction (c.f. Figure 3). However interesting these results are, we note that they could only be obtained when analyzing whole trial epochs, since limitations of the method – which requires long epoch lengths – hindered a spatially-resolved analysis.

**Figure 5.**
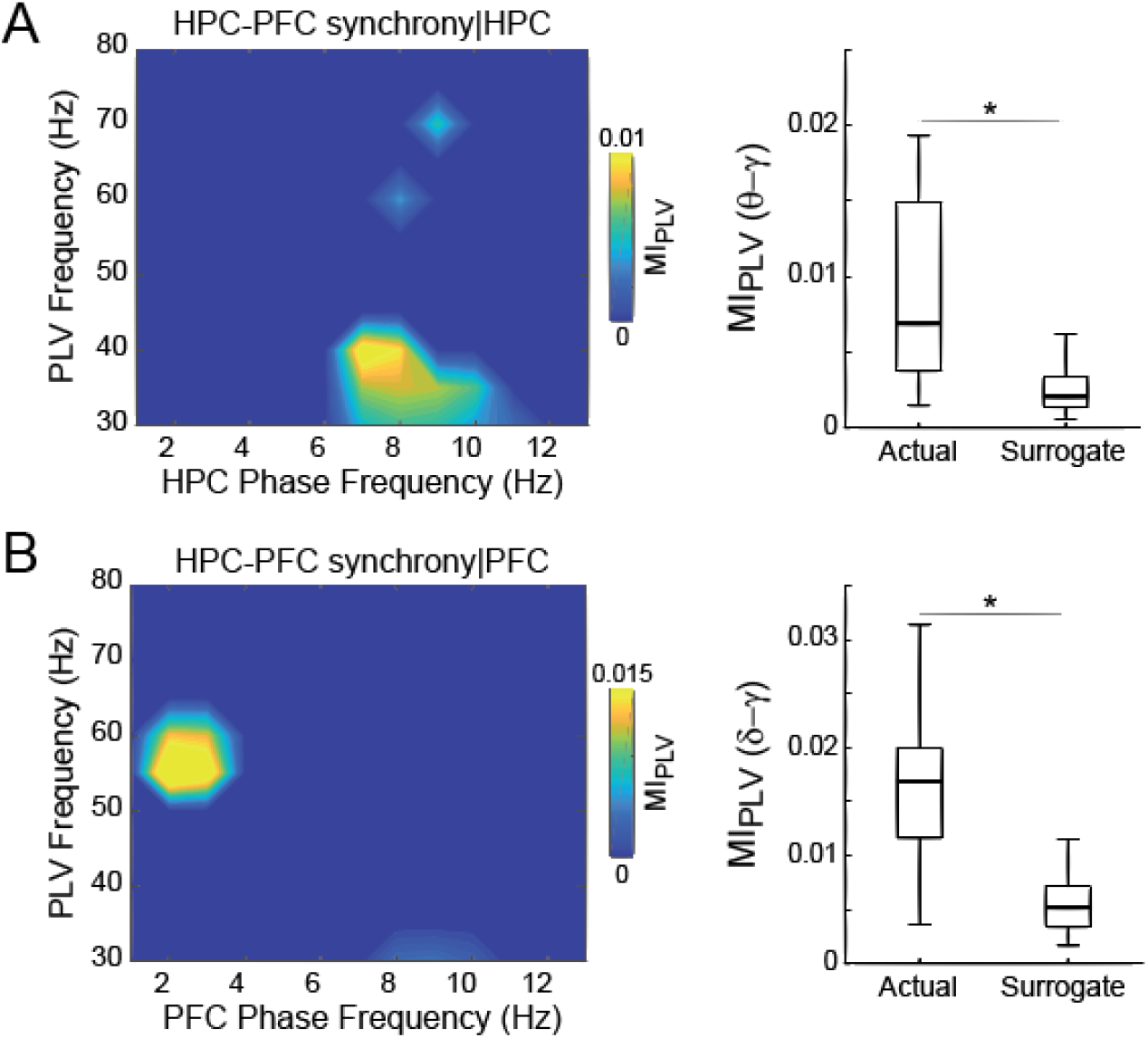
Hippocampal theta and prefrontal delta modulate inter-regional gamma synchrony at specific sub-bands. (A,B) (Left) Average mPFC-CA1 phase-synchrony comodulation map (n=13 sessions). The X-axis denotes the modulating phase frequency while the Y axis represents the fast frequencies analyzed for inter-regional synchrony, as measured by the phase-locking value (PLV). The color denotes the PLV modulation level for the fast frequency Y by the phase of the slow frequency X (MI_PLV_; see González et al., 2020). Only statistically significant MI_PLV_ values are shown (as assessed by a surrogate analysis, see Methods). The phase frequencies in A were obtained from the CA1 LFP, and in B from the mPFC. (Right) Boxplot distributions of actual (n = 13 sessions) and surrogate MI_PLV_ values for the relevant frequency pairs (HPC: 6-9 Hz vs 25-50 Hz; PFC: 1-4 Hz vs 45-70 Hz). *p<10^-30^, t-test.

Finally, in addition to the population-derived signal which is the LFP, we also examined neuronal spike data from both regions. We investigated local and inter-regional phase-locking of CA1 and mPFC action potentials to the LFP oscillations. As shown in Figure 6A, hippocampal neurons (n=442 units) exhibited higher phase-locking to the LFP theta phase than prefrontal ones (n= 1136 units; t(1576)=19.17, p<10^-73^, unpaired t-test). Moreover, the hippocampal theta oscillation modulated more strongly neurons in both CA1 and mPFC than the theta oscillation recorded in the mPFC (Figure 6A right; CA1 units: t(441)=19.23, p<10^-59^; mPFC units: t(1135)=9.60, p<10^-20^, paired t-tests).

**Figure 6.**
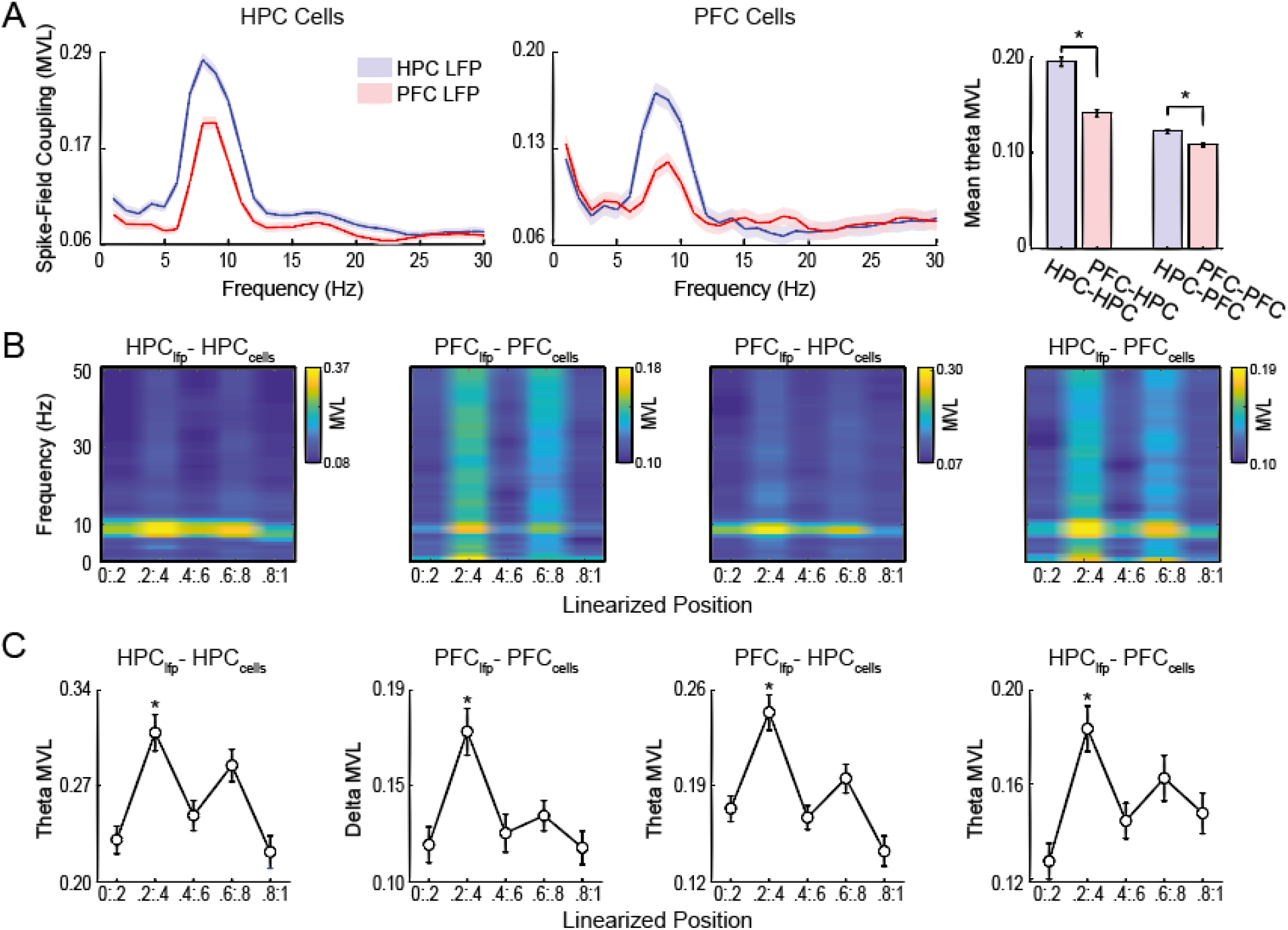
Prefrontal cortex neurons phase-lock to the hippocampal theta rhythm during maze runs. (A) Average spike-field coupling for local and inter-regional combinations (MVL: mean vector length). The hippocampal theta rhythm more strongly modulates neurons in both CA1 (n=442 units) and mPFC (n=1136 units). *p<10^-20^, paired t-test. (B) Spike-field coupling as a function of LFP frequency and space. (C) Mean spike-field coupling to theta or delta (for mPFC units) through space for significantly phase-locked cells (Rayleigh test; only cells with >10 spikes per spatial bin were taken into account, from left to right panels, n = 121, 82, 94 and 145 units). *p<10^-7^, repeated measures ANOVA.

Inspection of the spatial dynamics of intra- and inter-regional spike-field coupling corroborated the central role of the theta rhythm in modulating neuronal activity throughout the maze space in all regions (Figure 6B). In addition, mPFC neurons were also modulated by delta phase (Figure 6A,B); interestingly, the mPFC LFP exhibited a trend towards modulating its neurons more strongly at delta than theta (t(1135)=1.76, p=0.08, paired t-test). Hippocampal theta phase significantly modulated (Rayleigh’s circular test) 399 units in CA1 (90%) and 417 units in mPFC (37%), while prefrontal theta phase modulated 305 units in CA1 (69%) and 243 units in mPFC (21%). In addition, mPFC delta phase modulated 262 local units (23%). Figure 6C shows spike-field coupling strength across maze runs when only taking into account significantly phase-locked cells with >10 spikes per spatial bin. In all cases, the spike modulation was highest while animals were crossing the choice-related spatial bin (normalized positions from 0.2 to 0.4; p<10^-7^ for all cases under repeated measures ANOVA). Interestingly, during maze runs (normalized positions from 0.2 to 0.8), the local spike-field coupling strength of mPFC units at delta showed higher spatial selectivity for the choice-related spatial bin than the spike-field coupling of CA1 units at theta (mPFC spatial selectivity index: 0.39 ± 0.01 [M ± SEM]; CA1: 0.36 ± 0.01; t(201)=2.08, p=0.039, unpaired t-test). In all, these results are consistent with previous work examining the phase-locking of prefrontal neurons on similar tasks (Benchenane et al., 2010; Hyman et al., 2005; Jones and Wilson, 2005a).

## Discussion

Understanding how different regions of the brain communicate with each other, take part in solving tasks, and perform higher-level executive functions is a fundamental goal in neuroscience. Abnormal functioning of inter-regional connections is linked with various neuropathologies; in particular, activity in the prefrontal-hippocampal pathway can be seen as a marker for disorders such as Alzheimer’s disease and temporal lobe epilepsy when dysfunctional (Broggini et al., 2016; Kitchigina, 2018). The in vivo characterization of neural oscillations performed here can help to uncover the electrical signaling mechanisms of the brain and serve as a basis for further investigations. Our results point to specific oscillations such as delta (1-5 Hz) and theta (6-10 Hz) as possible means of communication between the hippocampus and prefrontal cortex. More importantly, some oscillatory features appear to be selective to task events associated with decision-making.

Our results show that the spectral power content of LFPs from both the hippocampus and prefrontal cortex varies as the animal progresses through the task. Namely, CA1 theta power increases during maze runs while PFC delta power has the opposite behavior (Figure 1B,C). But however interesting the spatial dynamics of theta and delta power may be, we could not confidently associate these power changes to putative differences in cognitive loads across task execution. Rather, the changes in power content were well matched by changes in locomotion speed. Moreover, when controlling for speed, we found no differences in theta nor in delta power for spatial regions before and after maze bifurcation (Figure 1D), which are typically assumed to be proxies for periods before and after the spatial decision (Jones and Wilson, 2005a; Benchenane et al., 2010). Nevertheless, the picture was different when we examined metrics evaluating the functional connectivity between the regions. Namely, we found that the inter-regional phase coherence at theta frequency was significantly higher before than after maze bifurcation (Figure 2), and this result held true even for spatial regions in which theta power was similarly high before and after the turn (Figure 2C).

The increase in hippocampal-prefrontal coherence observed here corroborates previous findings from rats subjected to spatial mazes, which pointed to a theta-frequency interplay as a mechanism underlying inter-regional communication and spatial decision-making. Among them, the pioneering work of Jones and Wilson (2005) showed that CA1-mPFC theta coherence is higher when rats traverse the central arm of the maze prior to performing a left-or-right spatial decision (“choice trials”) than during trials requiring no decision (“forced-turn trials”). Moreover, they also showed that theta coherence was higher for central arm runs subsequently associated with correct than error choices (Jones and Wilson, 2005). Five years later, Benchenane et al. (2010) showed a similar increase in theta coherence between LFPs from the hippocampus and mPFC at the choice point of a Y-maze task. Moreover, such a coherence increase was particularly prominent after animals learned the task rule (Benchenane et al., 2010). Similar increases in hippocampal-prefrontal coherence were also reported for mice recorded during spatial working memory tasks (Sigurdsson et al., 2010; O’Neill et al., 2013). Thus, in a time when scientific reproducibility has been put into question (Ioannidis, 2005; Collaboration, 2015), it is reassuring that the increase in CA1-mPFC theta coherence in rodents performing spatial decision tasks has been replicated by independent studies, including the present one.

Directionality is another major aspect that should be taken into account when investigating the interplay between brain regions. Here we computed frequency spectra of Granger causality (GC) (Granger, 1969; Geweke, 1984). These showed that the hippocampus has a causal influence on the prefrontal cortex at theta frequency and, interestingly, that the prefrontal-hippocampal causality is maximum at delta frequency (Figure 3). According to the operational definition, this means that the past values of hippocampal theta oscillations increase the prediction of the current value of the mPFC theta wave beyond what can already be predicted by its past values, and conversely for hippocampal delta activity. These results are consistent with the study by Zhan (2015), who also observed causality peaks at either theta or delta frequency depending on flow direction between these regions (e.g., Figure 7 in that study). But perhaps most strikingly, here we found that the magnitude of this directed functional connectivity varied across trial execution (Figure 3), suggesting that it could be related to different cognitive loads. Furthermore, HPC->PFC and PFC->HPC directionality peaked at different regions of the maze. The HPC->PFC GC peak early in the maze suggests that spatial information from the hippocampus is fed through theta oscillations into higher-order areas associated with executive functions at the beginning of the decision-making task, after the animal has left the initial box. On the other hand, the PFC->HPC GC peaked near the curve onset, suggesting that the prefrontal cortex sends feedback signals through delta oscillations shortly after the animal has committed to the spatial choice.

There is still a gap in our understanding of brain oscillations when it comes to the unification of local and inter-regional circuits. Local circuits tend to generate higher frequency oscillations such as gamma (Dickson et al., 2000; Buzsáki, 2006; Atallah and Scanziani, 2009), while lower frequencies, such as the theta and delta investigated here, tend to be associated with long-range communication (Astrid von Stein and Sarnthein, 2000; Hyman et al., 2005; Sirota et al., 2008). Cross-frequency coupling (CFC) is a suggested mechanism to act as an integrator between both sets of frequencies (Canolty and Knight, 2010). Based on this functional premise, here we have investigated the spatial dynamics of local and inter-regional phase-amplitude coupling (PAC) (Figure 4). Intra-regional PAC patterns were characterized by theta-gamma coupling in the hippocampus and delta-gamma coupling in the mPFC. Interestingly, and in accordance with previous findings (Zhong et al., 2017; Tort et al., 2018a), the modulated PFC gamma frequency was faster than the modulated CA1 gamma frequency, suggesting that these regions may use different gamma sub-bands for their local computations. Regarding cross-regional PAC, hippocampal gamma amplitude coupled to the mPFC theta, whereas mPFC gamma exhibited no meaningful coupling to hippocampal theta. While the latter finding contrasts with an influential previous study (Sirota et al., 2008), it corroborates a more recent study in mice that also reported no influence of hippocampal theta phase over the amplitude of prefrontal gamma (Zhang et al., 2016; see their Figure 6). Finally, we found that PAC strength varied as a function of space during maze runs (Figure 4). This finding is consistent with previous reports suggesting a role for CFC during spatial decision-making (Tort et al., 2008; Schomburg et al., 2014), and may reflect the computations required for different cognitive and behavioral demands across task execution.

Even though there were no simultaneous recordings of respiration in the analyzed dataset, a note about possible influences of nasal breathing over the observed findings is in order. In particular, since rats often breathe at delta frequency (Kay et al., 2009; Rojas-Líbano et al., 2014; Tort et al., 2018a) we believe that the prefrontal “delta” oscillations observed here (Figure 1) and in previous studies (Place et al., 2016; Andino-Pavlovsky et al., 2017; Guise and Shapiro, 2017) correspond to respiration-coupled oscillations, which were recently shown to be prominent in the mPFC (Biskamp et al., 2017; Zhong et al., 2017; Tort et al., 2018b). Furthermore, it should be noted that the pattern of delta-gamma coupling in the mPFC perfectly matches the pattern of respiration-gamma coupling previously described for this region (Tort et al., 2018a; Zhong et al., 2017). Interestingly, Place et al. (2016) reported theta causality at the HPC->PFC direction while animals explored an arena (consistent with our findings), but on the reverse direction (PFC->HPC) while animals sampled objects present in the arena. The latter finding apparently contrasts with ours, since here PFC->HPC causality occurred at delta, not theta (Figure 4). However, considering that the objects in Place et al (2016) were identical terra cotta pots with different odors, animals may have sniffed the pots during object sampling, giving rise to respiration-coupled oscillations at theta frequency (that is, at the breathing frequency during sniffing). Respiration-coupled oscillations, as theta, have also been suggested to mediate long-range interactions (Tort et al., 2018a). Whether they would be particularly more relevant during odor-guided decisions, as in the task investigated here, remains to be determined.

Using a new screening method (González et al., 2020), we were able to analyze the integration of two different mechanisms of long-range neural communication: CTC (Fries, 2015, 2005) and CFC (Canolty and Knight, 2010; Tort et al., 2008). That allowed us to discover a modulation of the inter-regional gamma synchrony by the phase of the slower oscillations, which moreover seemed to be sub-band specific (Figure 5). Namely, we found that the phase of mPFC delta modulates long-range 40-60 Hz synchrony, while the hippocampal theta phase modulates synchrony at the 30-40 Hz gamma sub-band. Unfortunately, we were not able to characterize the dynamics of this modulation along maze runs due to the method requiring long epoch lengths, hindering an analysis at the level of spatial bins. It remains to be determined whether these synchrony modulations are task-specific or not; at present, there are no other reports examining the phase-modulation of hippocampal-prefrontal CTC in other tasks given the recentness of the method.

We also analyzed spike data from both regions to investigate phase-locking dynamics locally and inter-regionally. Not surprisingly (Skaggs et al., 1996; Jensen, 2005; Foster and Wilson, 2007), hippocampal neurons showed higher levels of entrainment to theta oscillations than prefrontal neurons (Figure 6A), especially from its own LFP given their particular prominence (Winson, 1974; Rawlins et al., 1979; Buzsáki, 2002). Interestingly, PFC cells also showed higher phase-locking to the CA1 rhythm than to the locally recorded theta activity, despite the considerable distance between the regions. When characterizing the dynamics of spike-phase-locking through maze space, we again found a prominent role of theta in all local and cross-regional combinations of field potential and neuronal activity. In addition, high modulation by delta was also observed for the spike-field coupling within the mPFC (Figure 6B). These analyses further revealed that the magnitude of spike-field coupling varied across the maze, with maximal strength at the choice-related area (Figure 6C). Interestingly, the spatial dynamics of spike-field coupling within the mPFC exhibited a greater spatial selectivity for the choice area than within CA1, resembling the spatial profiles observed in our CFC analysis. These results are consistent with previous reports showing increased modulation of prefrontal cells by the hippocampal theta at the choice-related area in a variety of tasks (Hyman et al., 2005; Jones and Wilson, 2005b; Siapas et al., 2005; Colgin, 2011).

Our work has built upon previous studies on the hippocampal-prefrontal network interactions during decision-making that directly recorded from these regions (e.g., Jones and Wilson, 2005; Benchenane et al., 2010). However, even though there is by now abundant evidence showing a bidirectional functional link between these structures (Preston and Eichenbaum, 2013; Jin and Maren, 2015; Sigurdsson and Duvarci, 2016; Eichenbaum, 2017), it should be noted that the mPFC receives connections from the intermediate hippocampus (Swanson, 1981; Jay and Witter, 1991) but does not send direct projections back to it (Hoover and Vertes, 2007). Anatomical findings point to the thalamic nucleus reuniens as the likely candidate to mediate mPFC signals to the CA1 region (Vertes et al., 2007; Ito et al., 2015). Consistent with these reports, lesioning the nucleus reuniens in rats impairs performance in spatial memory tasks (Hembrook and Mair, 2011; Hembrook et al., 2012), especially those requiring route planning (Cholvin et al., 2013). Interestingly, recent electrophysiological studies have been providing functional evidence for the role of thalamic nuclei in mediating the communication between the hippocampus and the prefrontal cortex in animals performing spatial tasks (Ito et al., 2015, 2018).

In summary, we have here further characterized the electrophysiological signatures of the hippocampal-prefrontal interplay during spatial decision making. Our results add to others by showing dynamic patterns of electrophysiological interactions that can be related to the varying cognitive and behavioral demands of an odor-cued spatial alternation task. In particular, our results suggest that oscillations both at the network and spike levels constitute important mechanisms for inter-regional communication and that different frequency channels may be used depending on flow direction.

## Acknowledgements

This work was supported by the Brazilian Federal Agency for Support and Evaluation of Graduate Education (CAPES) and the Brazilian National Council of Technological and Scientific Development (CNPq). We thank the Buszáki laboratory for making the data set publicly available at the data sharing website crcns.org, Shigeyoshi Fujisawa for kindly providing additional data, Wilfredo Blanco Figuerola, Abner Rodrigues, César Rennó-Costa and Bryan Souza for thoughtful discussions.

## Methods

### Data and materials

The data used in this work was collected and kindly made available by the Buszáki Lab through the Collaborative Research in Computational Neuroscience data sharing website crcns.org (Fujisawa et al., 2015). The original descriptions for the surgical and experimental procedures can be found in Fujisawa et al. (2008). Part of their methods is reproduced below for convenience, followed by details of the data analysis performed here.

### Animals and behavioral task

Subjects were three adult male Long Evans rats (3 to 5 months old) performing an odor-cued spatial alternation task in a figure-eight T-maze, which contained a start area, a center arm, and left and right goal arms (Figure 1A). In the start area, animals were trained to nose-poke to receive an odor sample (chocolate or cheese) indicating the goal arm and associated reward to be found at its end (e.g., right for 300 mg of chocolate, left for 300 mg of cheese). The particular match between odor and arm varied across subjects. The animals underwent surgery after achieving a performance of >85% correct choices in five consecutive days. Position in the maze is tracked using two small light-emitting diodes mounted above the head stage and a digital video camera recording at 40 Hz.

### Data acquisition

Neuronal activity was recorded simultaneously from the medial prefrontal cortex with an 8-shank 64-channel probe and from the intermediate CA1 with a 4-shank 32-channel probe. Data was amplified 1000x and recorded at 20 kHz using a 128-channel DataMax system (16-bit resolution; RC Electronics). LFPs were obtained by low-pass filtering at 625 Hz and down-sampling to 1250 Hz.

### Spike detection and sorting

Data band-passed filtered from 0.8 to 5 kHz were used for spike detection. A mean RMS was computed using 100-s sections and a value of 7 standard deviations from the mean RMS was set as threshold. Spike sorting was done through PCA-derived features and semi-automatically clustered using KlustaKwik (Rossant et al., 2016).

### Data analysis

All analyses were performed with built-in and custom-written Matlab (Mathworks) routines.

### Spatial spectrograms and coherograms

Temporal spectrograms and coherograms were computed using the *spectrogram()* and *mscohere()* Matlab functions, respectively. For spectrograms, the window size was 500 ms with an overlap of 45%; for coherograms, we used 5-s windows with 97.5% overlap (i.e., step size = 125 ms). Within each window, the coherence spectrum was obtained using 1-s sub-windows with no overlap. Spatial spectrograms and coherograms were obtained as follows: first, for each trial we computed the temporal spectrogram or coherogram as described above; then, for each bin of space (see below), we average all spectrogram or coherogram columns (that is, power or coherence spectra) associated with the timestamps the animal spent in that spatial bin; finally, for each spatial bin we averaged all associated spectra across trials in a session. The spatial bins were defined by dividing the mean trajectory into 20 equal parts, called linearized positions, varying from 0 (start of the trial) to 1 (reward consumption). In addition to the average spectra associated to the spatial bins, in Figures 1B and 2A, we also show power and coherence spectra averaged for five 1-s time bins before and after maze runs.

### Task-event coherence

We analyze four unique events in the task: start, choice, turn and reward. The start event was defined as the timestamp of the nose poke in the initial box; the reward event as the timestamp when the reward started to be consumed; the choice and turn events were defined by the crossing of the linearized positions of 0.3 and 0.45, respectively. In Figure 2D, task-event coherence was computed using 500-ms time windows with 100-ms steps around the event timestamp. For each 500-ms window, coherence was computed across all trials in a session (that is, every trial gives rise to a phase difference vector, and coherence is obtained as the square of the length of the average vector). In terms of implementation, this can be achieved by concatenating the 500-ms windows across trials and employing *mscohere()* with 500-ms window length and no overlap. In Figure 2E, peak theta coherence was obtained as the mean peak value across the six peri-event windows (centered from −250 ms to +250 ms).

### Average speeds and speed controls

Instantaneous speed was computed as the animal displacement in the X-Y plane between two video frames (Euclidean distance) divided by the sampling interval of the camera (40 ms). The speed-controlled analysis shown in Figure 1D was performed by selecting only trials in which the average speed for the analyzed spatial sections was between 40 and 50 cm/s.

### Granger causality

Granger causality was computed using the Multivariate Granger Causality Matlab Toolbox (Barnett and Seth, 2014). To prevent oversized model orders and amplitude biases, we downsampled the LFP to 125 Hz and z-scored the signal. The Variational Autoregressive (VAR) model order was fixed as 15 for all analyses, estimated using the Bayesian Information Criterion (BIC). The task-event analysis shown in Figure 3A was computed using 500-ms windows centered on the task events concatenated across all trials in a session. The “spatial Grangerogram” in Figure 3B was obtained by computing Granger causality spectra for 10 equally-divided linearized position bins from trial start to trial end. To that end, for each position bin, the associated LFP epochs were first concatenated across trials.

### Cross-frequency coupling

Phase-amplitude coupling (Figure 4) is computed within and between regions using the Modulation Index method described in Tort et al. (2010) and available at https://github.com/tortlab/phase-amplitude-coupling. Filtering bandwidths were 4 Hz for phase frequencies (0.5 Hz step) and 10 Hz for amplitude frequencies (5 Hz step); the phase time series were binned into 18 equally spaced bins (i.e., 20° per phase bin). In Figure 4A, we computed phase-amplitude comodulograms using data from the whole session excluding the inter-trial periods. To obtain the phase-modulated spatial comodulograms shown in Figure 4B, we first computed 5 standard comodulograms, one for each position interval (a “spatial bin”). To that end, for each spatial bin, we concatenated the associated filtered LFP epochs across trials (i.e., to avoid edge effects, we first filter the whole signal to obtain the phase and amplitude time series, then we concatenate to compute the comodulogram; see Tort et al. 2008). We then averaged the comodulogram values over the phase frequency range of interest (4-6 Hz for PFC and 6-10 Hz for CA1). In Figure 4C, we further averaged MI values using the amplitude sub-band with the highest comodulation value, which varied across subjects (Rat EE: 60-70 Hz, Rat FF: 45-55 Hz, Rat GG: 65-75 Hz).

### Phase-synchrony comodulation

Phase-synchrony comodulation is a new screening method proposed by González et al. (2020) for detecting the modulation of gamma synchrony by the instantaneous phase of a slower oscillation. In short, the method uses a combination of the phase-locking index (PLV) and MI metrics to assess communication-trough-coherence (CTC, measured by the PLV) by means of cross-frequency coupling (CFC, measured by the MI, which in the context of this metric is referred to as MI_PLV_). Matlab codes for computing the MI_PLV_ are available at https://github.com/tortlab/Phase-Locking-Value---Modulation-Index. Here, slow- and fast-frequency LFP components were obtained by band-pass filtering with 2-Hz (1-Hz step) and 20-Hz (5-Hz step) bandwidths, respectively. The slow frequency phase was binned into 18 bins. We analyzed data from the whole session excluding the inter-trial periods, and, as in the CFC analysis, the signals were first filtered before extracting the instantaneous phases to avoid edge effects. Surrogate analysis was performed by randomly circular shifting the phase of slower oscillations between 1 to 10 seconds. This was done 100 times per session, resulting in a null hypothesis distribution of 1300 surrogate MI_PLV_ samples. To consider the synchrony modulation for a given frequency pair as statistically significant, the mean original MI_PLV_ of that pair (n = 13 sessions) had to be higher than the 95^th^ percentile of the surrogate values. In the phase-synchrony comodulograms of Figure 5A, the non-significant MIPLV values were set to zero.

### Spike-field coupling

The magnitude of spike-phase locking in Figure 6 was estimated by the mean vector length (MVL), also known as mean resultant length (MRL; Sigurdsson et al., 2010). The MVL is obtained by first representing each spike as a vector on the unit circle whose angle is the spike phase; then, the average vector (over all spikes) is computed and its norm (i.e, the MVL) determined. Phase-locked and non-phase-locked cells were defined using Rayleigh’s circular test for non-uniformity with α = 0.05. For the spatial analyses, we used 5 equally-divided linearized position bins from trial start to trial end: 0-0.2, 0.2-0.4, 0.4-0.6, 0.6-0.8 and 0.8-1. The epochs associated with each bin were concatenated across the entire session. To prevent imprecise MVL estimations due to low spike counts, we only included phase-locked cells with 10 or more spikes on each spatial bin. The spatial selectivity index for the choice-related spatial bin was defined as MVL_0.2-0.4_/(MVL_0.2-0.4_ + MVL_0.4-0.6_ + MVL_0.6-0.8_).

## Notes

### Competing Interest Statement

The authors have declared no competing interest.

